# The structure of genotype-phenotype maps makes fitness landscapes navigable

**DOI:** 10.1101/2021.10.11.463990

**Authors:** Sam F. Greenbury, Ard A. Louis, Sebastian E. Ahnert

**Affiliations:** Theory of Condensed Matter Group, Cavendish Laboratory, University of Cambridge, UK; NIHR Imperial Biomedical Research Centre, ITMAT Data Science Group, Imperial College London, UK; Department of Metabolism, Digestion and Reproduction, Imperial College London, UK; Rudolf Peierls Centre for Theoretical Physics, University of Oxford, UK; Department of Chemical Engineering and Biotechnology, University of Cambridge, Philippa Fawcett Drive, Cambridge CB3 0AS, United Kingdom; The Alan Turing Institute, British Library, 96 Euston Road, London NW1 2DB, United Kingdom

## Abstract

Fitness landscapes are often described in terms of ‘peaks’ and ‘valleys’, implying an intuitive low-dimensional landscape of the kind encountered in everyday experience. The space of genotypes, however, is extremely high-dimensional, which results in counter-intuitive properties of genotype-phenotype maps, such as the close proximity of one phenotype to many others. Here we investigate how common structural properties of high-dimensional genotype-phenotype maps, such as the presence of neutral networks, affect the navigability of fitness landscapes. For three biologically realistic genotype-phenotype map models—RNA secondary structure, protein tertiary structure and protein complexes—we find that, even under random fitness assignment, fitness maxima can be reached from almost any other phenotype without passing through a fitness valley. This in turn implies that true fitness valleys are very rare. By considering evolutionary simulations between pairs of real examples of functional RNA sequences, we show that accessible paths are also likely to be utilised under evolutionary dynamics.

## I. INTRODUCTION

Ever since they were first introduced in Sewall Wright’s foundational paper [1], fitness landscapes have become an enduring and central concept in evolutionary biology [2–6]. In particular, a low-dimensional picture of fitness ‘peaks’ and fitness ‘valleys’ has played an important role in shaping intuition around evolutionary dynamics. A key prediction is that a population must typically traverse an unfavourable valley of lower fitness to move from one fitness peak to another. But, as already pointed out by Fisher [7] and many others since [4, 8–11], the space of genotypes is typically extremely high dimensional. As illustrated in Fig. 1, what appears to be a fitness valley in a lower-dimensional landscape could be easily bypassed when dimensions are added [9–11].

**FIG. 1.**
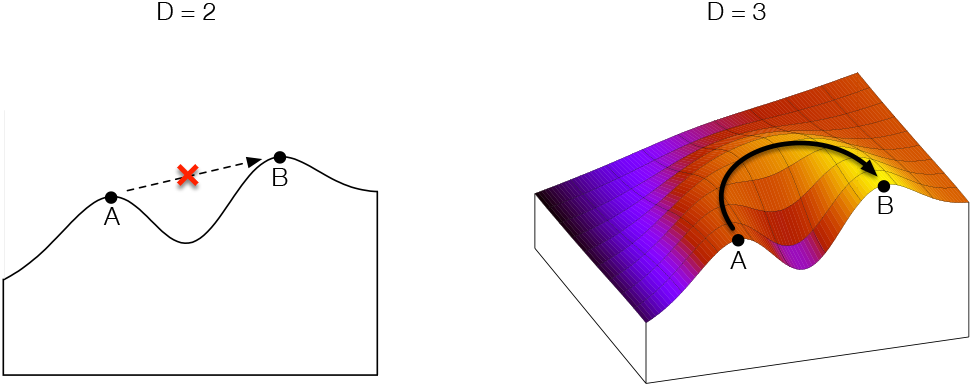
Illustration of how increasing dimensionality can affect the navigability and presence of valleys in a fitness landscape.

Two key open questions are: 1) Does the low-dimensional picture of fitness valleys hold for realistic high-dimensional genotype spaces? And 2), if we define *accessible paths* of point mutations between a low fitness phenotype and a high fitness phenotype as those with monotonically increasing fitness, are such paths sufficiently common that they can easily be found by an evolving population?

One way forward is to consider empirical fitness landscapes, where much recent progress has been made [5, 12], particularly for molecular phenotypes [5, 13–21]. This body of work has yielded important insights, such as the role of local epistatic interactions in sculpting evolutionary paths [22–24]. Nevertheless, ruling out high-dimensional bypasses is difficult in empirical studies because genotype spaces, which grow exponentially as *K*^*L*^ for alphabet size K and genotype length *L*, are almost always unimaginably vast [25]. They are also highly connected since distances are linear; two genotypes are at most *L* point mutations away, but are connected by up to *L*! possible paths. For example, even for a very short *L* = 20 strand of RNA, there are up to 20! ≈ 2× 10^18^ paths between any two genotypes. Empirical landscapes can typically only ever sample a small fraction of the full genotype space, so what may appear to be an isolated fitness peak, may in fact be accessible but the pathways are not feasible to experimentally identify.

A different strand of work, which can in principle address questions of global accessibility, has focused on model genotype-to-fitness landscapes [3, 6, 10, 11, 26, 27]. If fitness is assigned randomly to genotypes, as in Kingman’s ‘house of cards’ model [28], then the probability of finding accessible paths is small. If instead there are correlations between fitness and the genotypes, then, depending on details of the model, accessible paths can be common [11, 29]. While again much progress has been made in this literature, it is not always clear how well these models capture true biological fitness landscapes.

Here we take a different approach, and build upon recent advances showing that many realistic genotype-phenotype (GP) maps share key structural features that enhance navigability. [30–32]. One important commonality is the existence of large neutral networks of genotypes that map to the same phenotype. Because of these networks, the mutational robustness *ρ*_*p*_ of a phenotype *p* (defined as the mean probability that a point mutation leaves the phenotype unchanged) typically scales as the logarithm of phenotype frequency *f*_*p*_ (defined as fraction of genotype space occupied by phenotype *p*) for a wide range of GP maps [30–32]. If the genotypes of a phenotype were randomly distributed in genotype space, then the robustness would scale as *ρ*_*p*_ ≈ *f*_*p*_, which is much smaller than observed, highlighting the presence of neutral correlations in many realistic GP maps [30]. Large neutral networks play an important role in evolution because they allow all adjacent phenotypes of a neutral network to be reached by point mutations from any individual genotype in that network [31–34], and therefore may form part of accessible paths.

Our main contribution here is to show that commonly observed structural properties of GP maps greatly increase the number of accessible paths, or ‘navigability’, in associated fitness landscapes. In contrast to the genotype-to-fitness models studied by others (see above), we consider the genotypephenotype (GP) map with the phenotype-to-fitness map as an additional layer on top. We first explore specific features of GP maps that affect the navigability: redundancy (large neutral sets), frequency of the unfolded or trivial phenotype, neutral correlations and high-dimensionality, and the effect of these quantities on the ruggedness of the landscape. We then focus on identifying whether accessible paths exist for fRNA phenotypes identified *in vivo* from the fRNA database [35], and simulate evolutionary dynamics to explore whether accessible paths might be utilised in biological evolution. Our findings show that certain structural properties of GP maps give rise to navigable fitness landscapes, and that the resulting accessible paths are indeed likely to be exploited in the course of biological evolution.

## II. RESULTS

### A. Several well-studied genotype-phenotype maps induce navigable fitness landscapes

A wide range of different GP maps share common structural properties, including a much larger number of genotypes than phenotypes (redundancy), a heavily skewed distribution in the number of genotypes per phenotype (phenotype bias), and close proximity of genotypes belonging to the same phenotype (which can also be described in terms of positive neutral correlations or large phenotypic robustness) [30, 31]. Here we consider the RNA secondary structure GP map for sequences of lengths *L* = 12 and *L* = 15 (RNA12, RNA15) [36–42], the Polyomino lattice self-assembly GP map (*S*_2,8_, *S*_3,8_) [30, 43], and several HP lattice protein folding GP maps (two compact GP maps HP5×5 and HP3×3×3, and two non-compact ones HP20 and HP25) [44–46].

We performed computational experiments in which fitness is assigned to phenotypes randomly, and two phenotypes are chosen randomly from the set of all phenotypes as the ‘source’ and ‘target’.

The navigability ⟨*ψ*⟩ is defined as:

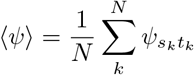

over a set of *N* source-target pairs (*s*_*k*_, *t*_*k*_), where *ψ*_*ij*_ is the probability that single-point mutation steps with monotonically increasing fitness (an accessible path) exist from a genotype of phenotype *i* to a genotype of phenotype *j*. In other words, the navigability is the average probability of an accessible path over the phenotypes of a GP map (see IV B 4).

In Table I, we report navigability for each GP map. The value of ⟨*ψ*⟩ is greater than 0.6 for all the GP maps we consider, apart from the non-compact HP models HP20 and HP25. The non-compact HP models have a navigability ⟨*ψ*⟩ ≤ 0.013 demonstrating these GP maps do not produce navigable fitness landscapes. These results suggest that the GP maps of RNA secondary structure, compact HP models, and the Polyomino model, have navigable fitness landscapes and contain very few fitness valleys under random fitness assignment. However, the lack of navigability in non-compact HP models highlights the need for further investigation of the effect of structural properties of the GP maps on navigability, which we pursue in the next section.

**TABLE I.**
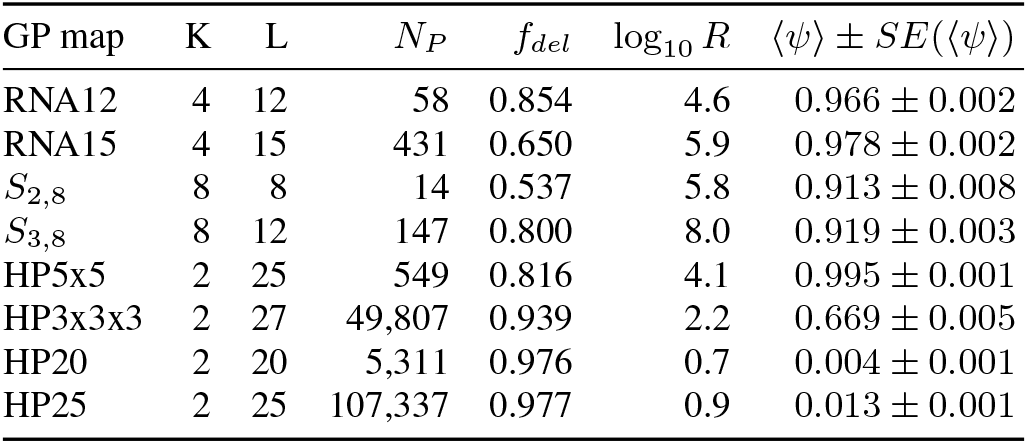
RNA, Polyomino and compact HP GP maps all have navigable fitness landscapes (⟨*ψ*⟩ > 0.6) under random fitness assignment illustrating a lack of fitness valleys. By contrast, non-compact HP models have very low navigability (⟨*ψ*⟩ ≤ 0.013).

### B. Common properties of GP maps are associated with navigability

#### 1. GP maps with fewer phenotypes and fewer deleterious genotypes are more navigable

Having showed that three distinct GP maps give rise to navigable fitness landscapes under random fitness assignment, we explore the relationship between structural properties GP maps and navigability. Specifically, we consider the redundancy *R* of a GP map, measured as the average number of genotypes per non-deleterious phenotype (see Eq. (1)), and the deleterious frequency *f*_*del*_. The deleterious frequency describes the fraction of genotype space that does not map to a well-defined phenotype. In the case of RNA secondary structure the deleterious phenotype would correspond to the unfolded RNA strand (i.e. the absence of any secondary structure). In the HP model it corresponds to the absence of a unique folded ground state. In the Polyomino model it corresponds to unbounded or non-deterministic assembly. In Fig. 2A we plot navigability against redundancy, while in Fig. 2B navigability is shown against the deleterious frequency with the numerical values provided in Table I. We observe a general increase in navigability for greater redundancy and smaller *f*_*del*_. HP3×3×3 presents an example of particular interest by maintaining navigability (⟨*ψ*⟩ = 0.669) with less redundancy (log_10_ *R* = 2.2) and large deleterious frequency (*f*_*del*_ = 0.939).

**FIG. 2.**
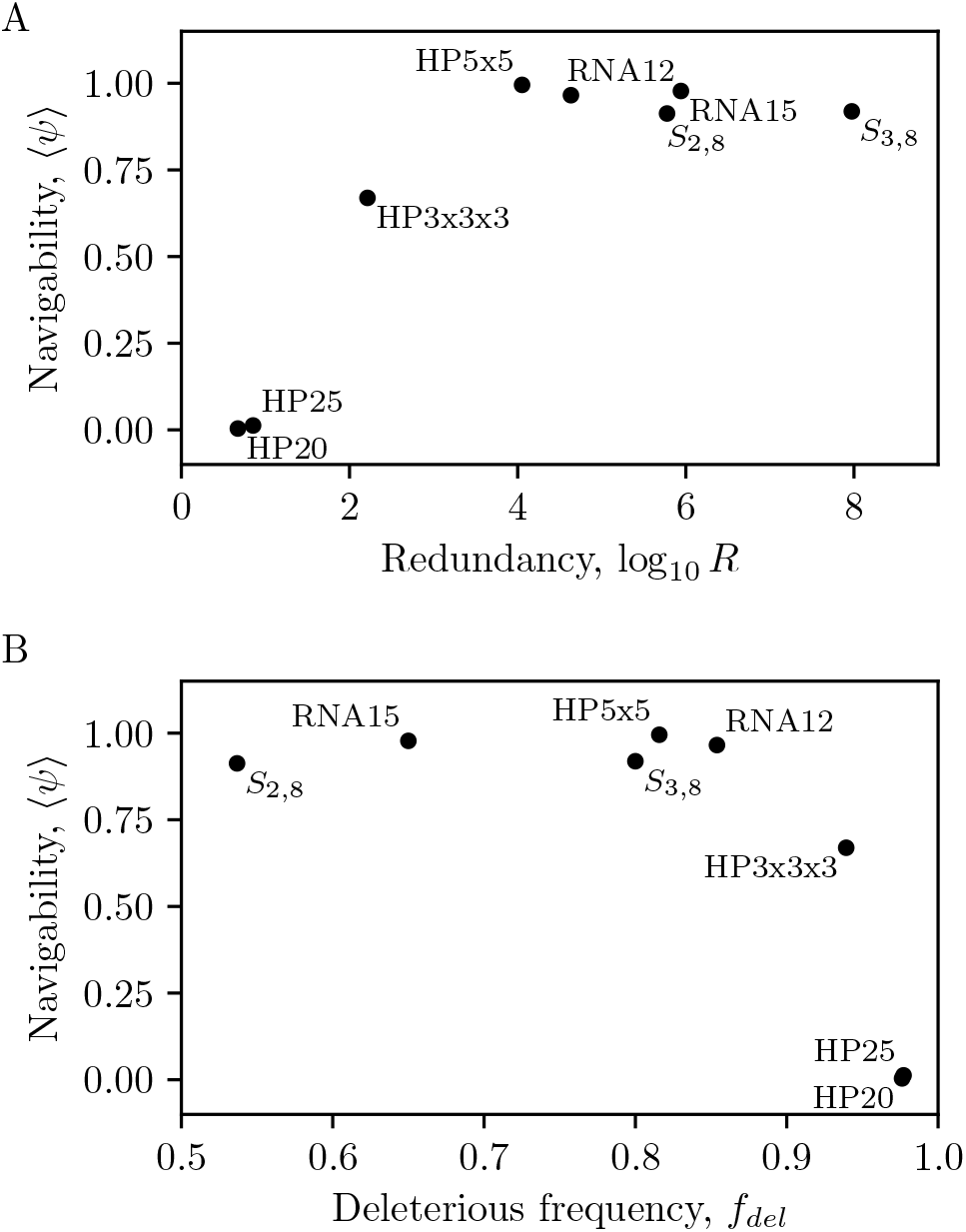
Navigability of each GP map is plotted in relation to (A) redundancy log_10_ *R* and (B) deleterious frequency *f*_*del*_. We find that there is a positive association with navigability and redundancy while a negative association with respect to deleterious frequency for large *f*_*del*_.

The results across different GP maps provide some intuition for factors that determine navigability. With decreasing redundancy, it becomes more difficult to access all phenotypes as they begin to occupy smaller fractions of the overall space. As *f*_*del*_ increases, more neighbours of a given genotype will have a fitness of 0, therefore localising phenotypes to smaller components in the GP map, increasing the likelihood of each genotype having no neighbouring genotypes with greater fitness.

#### 2. Positive neutral correlations increase navigability

We next consider how neutral correlations, which fundamentally arise from a very general picture of constrained and unconstrained portions of genotype sequences [47–49] and lead to greatly enhanced mutational robustness [30], affect navigability. The level of correlations in a given GP map can be adjusted by taking two genotypes *g*_1_ and *g*_2_ at random and assigning the phenotype of *g*_1_ to *g*_2_ and vice versa. Such random swaps remove the local correlations that are intrinsic to the GP map. The total number of swaps applied is parameterised as *s*. With increasing *s*, we decorrelate the GP map towards a random phenotype assignment to the set of genotypes. While *s* parameterises the decorrelation process, it is not on a scale that captures the level of correlations present relative to either the original GP map or fully randomised GP map where there are no correlations. Therefore, a measure of correlations *c*(*s*) (Eq. (9)) after *s* swaps is captured by relating phenotype robustness *ρ*_*p*_ and frequency *f*_*p*_ averaged across the phenotypes of the GP map for a given number of swaps *s*. When *c*(*s*) = 1, the correlations are equal to the original GP map, when *c*(*s*) = 0, the correlations are that of the randomised null model. Positive neutral correlations are present for *c*(*s*) > 0. By measuring the navigability ⟨*ψ*⟩ after a given number of swaps *s*, we measure the extent to which neutral correlations *c*(*s*) affect navigability.

In Fig. 3A, we plot how navigability varies with *c*(*s*) in *S*_2,8_, RNA12, HP5×5 and HP3×3×3 GP maps, a subset of the GP maps in the previous section that are both small enough to be tractable here, and have sufficiently large navigability such that the effect of reducing correlations and dimensionality may be sizeable. All four GP maps, on average, show greater navigability for greater *c*(*s*) with an approximately linear decay in navigability with decreasing *c*(*s*), saturating at a lower value specific to each GP map: 0.378 ± 0.005 for RNA12, 0.100 ± 0.003 for HP5×5, 0.000 ± 0.000 for HP3×3×3, and 0.949 ± 0.002 for *S*_2,8_, substantial reductions apart from for *S*_2,8_. In *S*_2,8_, the navigability ⟨*ψ*⟩ takes a greater value for the decorrelated GP map (*c* < 1) than for the original one (*c* = 1). This is because not all phenotypes are directly accessible from each other in the original GP map. However, a slight randomisation increases phenotype inter-connectivity due to the fact that the number of phenotypes for *S*_2,8_ is smaller than the number of local mutations (*N*_*P*_ < (*K* – 1)*L*). We expect that in GP maps of longer sequence length *L*, the role of positive neutral correlations will become even more pronounced. We explore this in Section II C with respect to fRNA phenotypes.

**FIG. 3.**
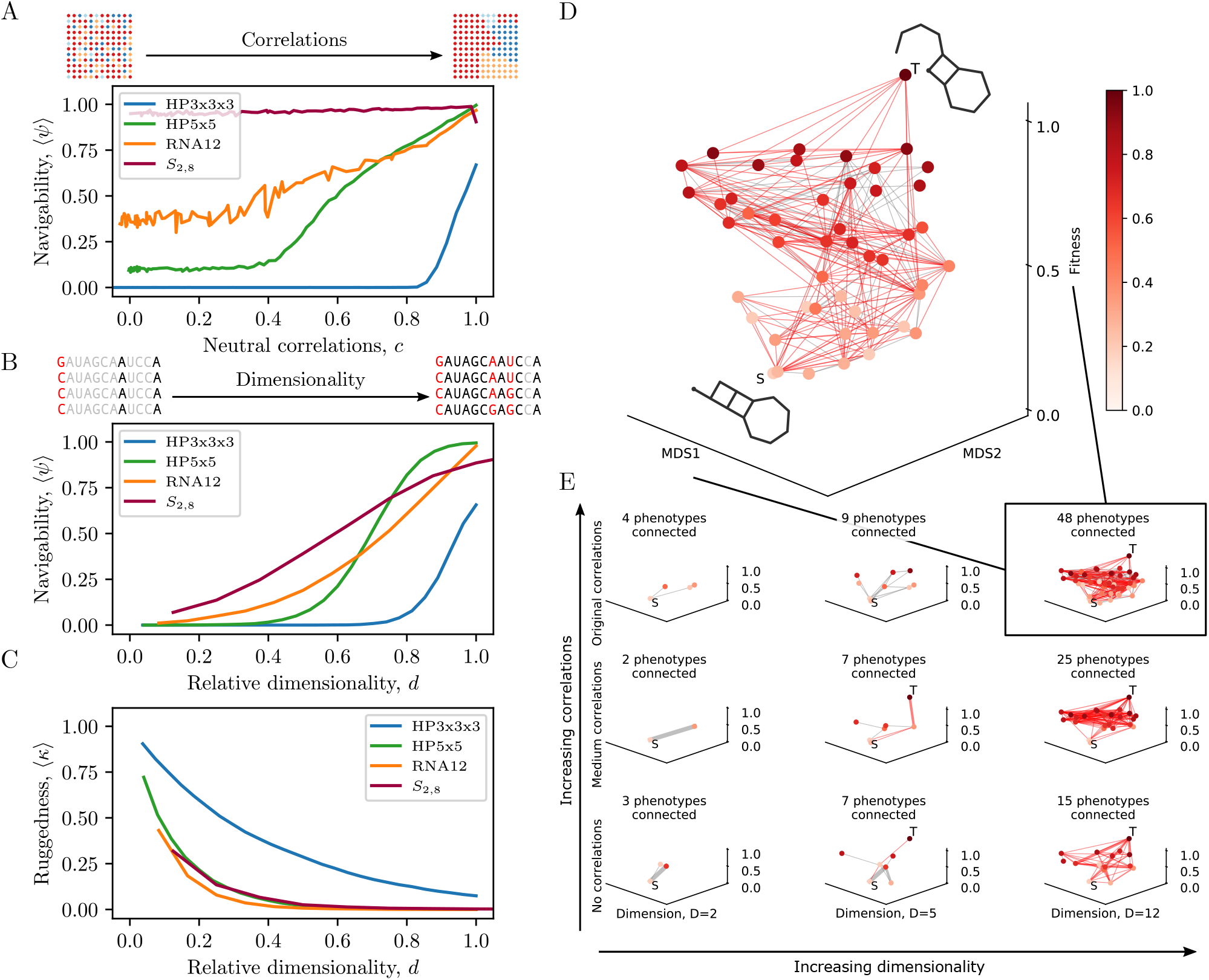
**(A) Navigability emerges as positive neutral correlations are added to HP3×3×3, HP5×5, RNA12 and *S*_2,8_ GP maps.** The level of neutral correlations is adjusted through genotype swaps, and the extent of positive neutral correlations after *s* swaps is measured on a scale *c* between the original GP map (*c* = 1) and the random null model’s correlations (*c* = 0). A caricature of the genotype space, coloured according to phenotypes, is shown for low neutral correlations (top left) and high neutral correlations (top right). **(B) Greater dimensionality of the GP map increases navigability for *S*_2,8_, HP3×3×3, HP5×5 and RNA12 GP maps.** During the search from a randomly chosen source phenotype to a target phenotype, we only allow *D* (*d* = *D/L*) of the total *L* bases to be mutated to explore genotype space. A caricature of a sequence with grey bases (*L* – *D*) not mutable, black bases mutable (*D*) and red bases varying across sequences, is depicted for low dimensionality (top left, *d* = 3/12) and high dimensionality (top right, *d* = 11/12). The GP maps show differing tolerance with respect to navigability under a change in dimensionality, *S*_2,8_ permitting navigability for low dimensionality significantly more than HP3×3×3, for example. (**C**) **With increasing dimensionality, landscape ruggedness decreases.** We measure landscape ruggedness ⟨*κ*⟩ as the average proportion of all genotypes encountered that are local fitness maxima (no neutral neighbours or neighbours with increased fitness). Ruggedness decreases in all GP maps as dimensionality increases, but the level of ruggedness is GP map dependent. **(D) A schematic of the joint effect of dimensionality and correlations on navigability through visualisation of the phenotype connectivity network.** An example is illustrated of the search for an accessible path in a specific random instance of a fitness landscape with the phenotype network of RNA12. Phenotypes are nodes and the edges are possible transitions between genotypes of those phenotypes given the random fitness assignments. Edges that are red are transitions that may lead to the target phenotype from the source phenotype. Inaccessible transitions are shown in grey. The vertical axis is fitness. The horizontal plane is a two-dimensional embedding of the phenotype space of RNA12 derived through a multidimensional scaling (MDS) that uses the pairwise Hamming distances between the dot-bracket representations of the phenotypes. It follows that proximity in the horizontal plane corresponds to similar dot-bracket phenotypes. **(E) The phenotype network is shown for three levels of correlations (original, medium, and no correlations) and three levels of dimensionality (*D* = 2, 6, 12).** Navigability and connectivity in the phenotypic network visibly increases with both increasing correlations and dimensionality.

#### 3. Large dimensionality increases navigability and decreases ruggedness

We now examine the effect of *dimensionality* of the GP map. The dimensionality of the entire GP map is defined as *L*, the length of the sequence. During the search for an accessible path from the source to target phenotype, all bases can be mutated, making use of the full dimensionality of the GP map. We can, however, reduce the dimensionality of the search by allowing only a random set of *D* sites (where *D* < *L*) to be mutated during a given search for an accessible path from source to target. We then consider ⟨*ψ*⟩ as a function of the *relative dimensionality* d = D/L ∀D ∈ {1, …, L}.

In Fig. 3B, we plot navigability ⟨*ψ*⟩ as a function of *d*. Reduced dimensionality severely reduces the navigability of fitness landscapes, with a sigmoidal relationship between ⟨*ψ*⟩ and *d*. All the curves show an increase from low navigability to high navigability as *d* → 1 of the full GP map. The critical value of *d*, and general scale and shape, is different across the four GP maps indicating a complex dependence on other GP map properties.

In addition to identifying an accessible path during the search from source to target, we also count the number of genotypes that do not have a neutral neighbour or neighbour with greater fitness. In other words, the proportion of genotypes that are local fitness peaks, therefore providing a measure of landscape *ruggedness*. The average proportion of genotypes that are local fitness peaks across source-target phenotype pairs and fitness assignments in a given GP map, is represented as ⟨*κ*⟩. In Fig. 3C, the ruggedness for each relative dimensionality *d* = *D/L* is plotted in the same four GP maps. We observe increasing dimensionality reduces ruggedness and, as relative dimensionality drops below a certain level, ruggedness sharply increases. Of note is HP3×3×3, where ruggedness is greater at a given relative dimensionality than for the other GP maps. Where all bases may mutate at *d* = 1, around 7 in 100 genotypes are local peaks (⟨*κ*⟩ = 0.07) but navigability remains high (⟨*ψ*⟩ = 0.66), demonstrating that partially rugged landscapes can still be navigable.

We illustrate an example of a source-target search in a schematic of the RNA12 GP map in Fig. 3D. We choose a random source and target pair and, during the search for an accessible path, keep track of all phenotypes encountered, their fitness and any transition between phenotypes. Each phenotype is represented as a node, edges as transitions between phenotypes, and the value on the vertical axis as the fitness. The *N*_*P*_ = 58 phenotypes of this GP map are assigned coordinates in the horizontal plane using multidimensional scaling (MDS) based on the pairwise Hamming distance between phenotypes [50]. This allows phenotypes that are similar to each other to be located in similar parts of the MDS1-MDS2 plane. The source and target phenotypes are labelled S and T respectively, edges that may form accessible paths are coloured red, and the remaining edges grey. This depiction of the fitness landscape immediately shows that it is highly connected with many accessible paths.

In Fig. 3E, with the same schematic source-target pair and fitness assignments as Fig. 3D, we illustrate the joint effect of neutral correlations and dimensionality on connectivity and navigability. We show the navigability of the phenotype network for three different degrees of correlation (no correlations, some correlations, original correlations) and three different dimensionalities (*D* = 2, 6, 12). The top right of the 9 plots is the original GP map that is also shown enlarged in Fig. 3D. We observe that decreasing both correlations and dimensionality of the search visibly reduces the navigability of the landscape through increasingly restricted networks. In the case of *D* = 2, the dimensionality in which fitness valleys are often visualised in the literature, phenotypic connectivity is sparse, making the landscape unnavigable. The increase in navigability with increases in both dimensionality and correlations highlight that both the structure of the underlying GP map and the high-dimensional nature of the evolutionary search are essential for fitness landscapes to be navigable.

### C. Navigability of functional RNA fitness landscapes

Next we focus on the RNA secondary structure GP map by specifically choosing source and target phenotypes that have been observed in nature. This is important as only a small subset of all possible phenotypes are typically seen in real biological systems [51, 52] and it is navigability among this subset that has most relevance for evolutionary processes.

#### 1. Fitness valleys are not observed between short fRNAs

We sample RNA secondary structures from the functional RNA database (fRNAdb) [35]. We consider pairs of fRNA phenotypes from the database with a given sequence length *L*, assigning a random fitness *F*_source_ ∈ [0, 1) and *F*_target_ = 1, with random uniform assignment of fitness for all non-trivial phenotypes found during the search process. We consider larger *L* than earlier, specifically in the range *L* ∈ [20, 40]. We perform two distinct types of search by either permitting or preventing neutral mutations in exploring a given genotype’s mutational neighbourhood. This provides a means to directly measure the role of neutral correlations in facilitating navigability for larger *L*. As the sequence length increases the number of phenotypes grows as *N*_*P*_ ≈ 1.76^*L*^ [53] producing a large computational overhead to track all phenotypes encountered during a search. In Section IV F, we describe in detail the more complex approach taken to measure navigability for larger *L*, which is necessary due to the increased computational expense.

In Table II, the navigability ⟨*ψ*⟩ for fitness landscapes with fRNA of sequence length *L* = 20 – 40 is reported along with the proportion of searches that were aborted and whether or not neutral mutations were permitted. With neutral mutations allowed, navigability is almost always 1.0, suggesting that fitness landscapes with fRNAdb source and targets are highly navigable. For *L* > 30 the proportion of aborted searches increases, leading to the greater potential for this estimate to be biased. However, there is a strong indication that with a greater computational threshold, similarly large navigability would be achieved at even larger *L* fRNA landscapes due to the observed scaling of ⟨*ψ*⟩ with the computational threshold (see Section A).

**TABLE II.** The navigability ⟨*ψ*⟩ for length *L* = 20 – 40 fRNAs, the number of unique targets tested, the number of phenotypes in the fRNA database, the proportion of runs that are aborted and the estimated navigability. Results for simulations with and without neutral mutations are shown in the left-hand and right-hand sets of columns respectively. For non-aborted runs with neutral mutations permitted, random observed fRNA landscapes are almost completely navigable. When neutral mutations are prohibited, navigability is severely reduced, but still substantial.

| L | Neutral mutations |  |  |  | No neutral mutations |  |  |  |
| --- | --- | --- | --- | --- | --- | --- | --- | --- |
| | Targets | $N_P^{\text{fRNAdb}}$ | $\langle\psi\rangle$ | Aborted | Targets | $N_P^{\text{fRNAdb}}$ | $\langle\psi\rangle$ | Aborted |
| 20 | 50 | 14350 | $0.987 \pm 0.004$ | 0.002 | 50 | 14350 | $0.530 \pm 0.016$ | 0.001 |
| 25 | 49 | 12958 | $0.999 \pm 0.001$ | 0.015 | 49 | 12958 | $0.661 \pm 0.015$ | 0.001 |
| 30 | 50 | 42195 | $1.000 \pm 0.000$ | 0.065 | 50 | 42195 | $0.638 \pm 0.015$ | 0.001 |
| 35 | 46 | 752 | $1.000 \pm 0.000$ | 0.190 | 46 | 752 | $0.599 \pm 0.016$ | 0.001 |
| 40 | 47 | 662 | $1.000 \pm 0.000$ | 0.517 | 47 | 662 | $0.380 \pm 0.015$ | 0.001 |

Where neutral mutations are disallowed, we find that navigability is markedly reduced below 1.0, although still substantially greater than zero (⟨*ψ*⟩ ∈ [0.38, 0.64]). The proportion of aborted searches is negligible. This finding is intriguing as it highlights that positive neutral correlations are important, but not essential, for the existence of accessible paths. A possible explanation lies in the vast number of phenotypes *N*_*P*_ ≈ 1.76^*L*^ available in the GP map, coupled with its high dimensionality. As fitness is randomly assigned and novel variation is only a few mutations away, there is a pool of non-neutral phenotypes with possibly larger fitness, potentially within a small mutational radius.

In Fig. 4, we use the representation introduced in Fig. 3D to illustrate an accessible path in fRNA. For the successful traversal between a specific source and target fRNA, we see a vast array of background, ‘greyed out’ phenotypes discovered during the search for an accessible path, as well as a shortest accessible path connecting 10 different phenotypes with the node colour and their vertical axis coordinate showing their fitness. This illustration further highlights the hyper-connectedness and high-dimensional bypasses present in fRNA GP maps that are afforded through exponentially increasing redundancy, positive neutral correlations, and high dimensionality. The phenotype network also serves again as an alternative depiction of the fitness landscape in which the effect of GP map structure on the course of potential evolutionary explorations may be grasped more intuitively.

**FIG. 4.**
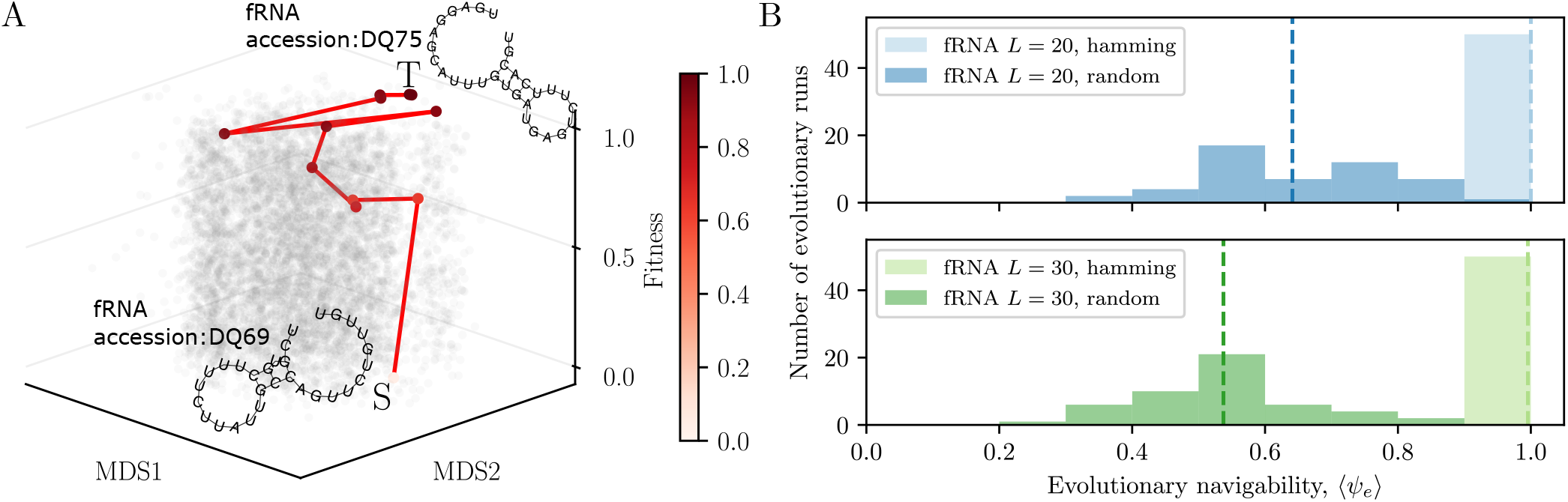
**(A) Example of an accessible path for a specific *L* = 30 fRNA source-target pair.** As introduced in Fig. 3D, phenotypes are nodes whose coordinates are derived from a multidimensional scaling (MDS) embedding of the phenotype similarities based on Hamming distance, while the vertical axis is fitness. We show the vast extent of phenotypes discovered during the search as ‘grey’ nodes, a shortest accessible path connecting the source and target phenotypes with red edges, and the phenotypes along this path shaded in proportion to fitness. The example illustrates the interconnected nature of the fitness landscapes for a concrete fRNA example, where the properties of the GP map are key in facilitating navigability. **(B) Evolutionary dynamics for fRNA with the distribution of *ψ* for randomly chosen phenotypes that belonged to the fRNA database were considered for the fixed *L* = 20 and *L* = 30 GP maps.** The navigability for *L* = 20 and *L* = 30 fRNA for 50 different target fRNA phenotypes are illustrated using histograms of ⟨*ψ*_*e*_⟩ for each target phenotype. The dark shaded bars show the proportion of successful searches for random fitness assignment, and the light bars for Hamming fitness assignment. Mean navigability of ⟨*ψ*_*e*_⟩ > 0.5 is observed for random fitness assignment and ⟨*ψ*_*e*_⟩ > 0.9 for Hamming fitness assignment.

Summarising our results, we have demonstrated that fRNA GP maps have navigable fitness landscapes up to *L* = 30 fRNA. They are highly likely to be navigable for even larger *in vivo* fRNAs due to the observed scaling of both the GP map properties and navigability with respect to the computational threshold. Neutral mutations drastically increase navigability but do not solely determine the presence of accessible paths.

### D. Evolutionary dynamics make use of accessible paths between fRNAs

Having considered whether accessible paths *exist* in a variety of GP maps, we consider whether these accessible paths are *utilised* under evolutionary dynamics.

It is conceivable that, while accessible paths exist in a fitness landscape, they may not be frequently used due to the entropic effects associated with the evolutionary search process. For example, if there are many mutational paths that lead to a local fitness maximum compared to a single path leading to the globally fittest phenotype, the increased number of ways to reach the local peak may result in populations taking one of these more prevalent paths and becoming trapped, necessitating passage across a fitness valley to reach the fittest phenotype.

We simulated evolutionary dynamics with a Wright-Fisher process, implemented via a genetic algorithm, and considered two different fitness assignment schemes: (a) random and (b) using a given phenotype’s dot-bracket Hamming distance to the target phenotype.

We chose *N*_*s*_ = 50 source phenotypes for each of *N*_*t*_ = 20 target phenotypes. During an evolutionary search, the change in fitness of the majority phenotype in the population was measured. The population was initialised from a population of genotypes that map to the source phenotype, and the fitness of the target was set to 1. We consider only the set of evolutionary simulations where the population was able to reach the target phenotype. We define *evolutionary navigability* ⟨*ψ*_*e*_⟩ as the average probability that the population’s majority phenotype reaches a target phenotype from a source phenotype (both randomly chosen) via an accessible path.

We consider only the polymorphic dynamical regime (*NμL* ≫ 1, where *N* is population size, *μ* is point mutation rate and *L* is genotype length). This case provides dynamics that are most likely to be associated with entropic regimes due to rapid discovery and exploration of mutational pathways that lead to more prevalent local fitness peaks as opposed to global ones. A greater mutation rate also increases the ability to cross fitness valleys making it a valuable test under which to consider whether evolutionary accessible paths continue to be used. Further details of the evolutionary simulation are provided in the methods (see Section IV G).

In Fig. 4B the navigability for *L* = 20 and *L* = 30 fRNA is illustrated with histograms binning the value of ⟨*ψ*_*e*_⟩ for each of *N*_*t*_ = 50 target phenotypes. The darker-shaded bars show the proportion of successful searches for random fitness assignment, and the lighter-shaded bars for the fitness assignment based on dot-bracket Hamming distance from the target phenotype. The mean navigability ⟨*ψ*_*e*_⟩ is shown as a vertical dashed line. For random fitness assignment we find decreased navigability values of ⟨*ψ*_*e*_⟩ ≈ 0.64 for *L* = 20 and ⟨*ψ*_*e*_⟩ ≈ 0.54 for *L* = 30 compared to the non-evolutionary scenario, for which ⟨*ψ*⟩ ≈ 1. While this is a reduction relative to the potential navigability present in the landscape, this still suggests that accessible paths are utilised for the majority of targets.

Under Hamming distance fitness assignment we found that accessible paths are taken much more frequently with all target phenotypes having ⟨*ψ*_*e*_⟩ = 1.0 for *L* = 20 and ⟨*ψ*_*e*_⟩ > 0.94 for *L* = 30. This provides additional evidence of evolutionary navigability with a plausible alternative fitness assignment and identifies the potential importance of *phenotypic correlations* within the GP map (in addition to the genotypic correlations discussed above) for evolutionary navigability.

## III. SUMMARY AND DISCUSSION

In this paper, we considered the navigability of fitness landscapes and, specifically, whether fitness valleys are prevalent in high-dimensional fitness landscapes based on biologically realistic GP maps. We examined three such GP maps with common structural properties and found that they were highly navigable, suggesting that fitness valleys are largely absent. We generalised this by demonstrating navigability in GP maps with longer RNA sequences using phenotypes contained in the fRNAdb database of fRNA observed in nature. Finally, we considered the question of whether accessible paths not only exist, but are also utilised by populations subject to evolutionary dynamics. We found that accessible paths are followed frequently by populations in evolutionary simulations, and are therefore likely to play an important role in real evolutionary settings.

We identified that universal structural properties of GP maps can facilitate navigability, namely: genotypic redundancy, the frequency of the deleterious phenotype, positive neutral correlations, and high dimensionality as a proportion of sequence length. These are important factors that are not characterised in a direct genotype-to-fitness mapping and are necessary to provide navigability. Additionally, we demonstrated that the phenotype network is arguably a more useful way to conceptualise evolutionary exploration. Visualising the fitness landscape in this way avoids the misleading intuitions of fitness valleys that can arise from the low-dimensional fitness landscape metaphor.

While we found fitness landscapes to be generally navigable under evolutionary dynamics, this navigability was lower than one might expect given the potential availability of accessible paths in these landscapes. We suggest two possible reasons for the reduction in the evolutionary setting: 1) the population truly arrives at a local optima and has no choice but to cross a fitness valley, and 2) due to the stochastic nature of the evolutionary dynamics the fitness of the population’s majority phenotype may drop, but not all members will necessarily have a reduced fitness and therefore, for the majority to return to a greater fitness, a fitness valley may not need to be crossed. This may lead to an underestimate of the true evolutionary navigability. The sensitivity of evolutionary navigability to alternative definitions is an important area for future exploration. The replacement of the random fitness assignment with one based on Hamming distance improved navigability drastically and highlights the role that *phenotypic correlations* play in GP maps in addition to the genotypic correlations discussed in [30].

A central assumption was that function and fitness are directly related to shape of the physical structure alone. This is an assumption made ubiquitously in the study of self-assembly GP maps where the structure is the sole component of the phenotype [31, 32]. Importantly, this will not always hold for all biological systems. For example, where a specific sequence is necessary to facilitate binding of a protein, an additional sequence constraint is imposed on top of that required to specify the structure. This additional specificity potentially reduces both the redundancy of the phenotype and the dimensionality available for accessing alternate genotypes. Our findings regarding landscape navigability should therefore be considered in the context of the GP map properties that facilitate accessible paths. If these properties are not present in the system in question, the fitness landscape is unlikely to be navigable. This is supported, for example, in ref. [54] where fragmented fitness peaks are identified in a rare exhaustive empirical fitness landscape study, but where fitness was specifically determined by the ability for GTP to bind rather than by specific secondary structure itself.

The metaphor of the fitness landscape has endured for almost a century of research in evolutionary biology. It is often discussed in intuitive terms, as a low-dimensional landscape. This can be problematic, as it obscures counter-intuitive properties of high-dimensional spaces, which real fitness landscapes are. Moreover, much of the literature on fitness landscapes does not consider genotype-phenotype maps and their properties, such as the ubiquity of neutral networks and their correlations in genotype space. Our contribution demonstrates that specific GP map properties, in combination with high-dimensionality, make fitness landscapes navigable. We show that accessible paths are not only available in three different biologically realistic GP maps, but also that they are followed in simulated evolutionary dynamics of functional RNA structures. These findings demonstrate that fitness valleys are largely absent in three biological GP maps. Given that the relevant GP map properties have been found in numerous other GP maps, it is highly likely that fitness valleys are indeed uncommon across a wide range of biological systems. Our findings support work on the role of high-dimensionality in promoting accessibility [11], as well as attempts to create an up-to-date metaphor for evolutionary adaptation [55]. A fuller understanding of the role of the GP map in structuring the high dimensional fitness landscape could provide vital insights into areas such as the arrival of drug resistance [56, 57] or the mutational progressions of cancer [58].

## IV. METHODS

### A. Self-assembly GP maps

We consider three GP maps for different systems of biological self-assembly: the RNA secondary structure GP map for secondary structure of RNA sequences, the HP lattice model for protein tertiary structure [44, 59] and the Polyomino model for protein quaternary structure [43]. The phenotype in each is solely related to the assembled structure. We briefly summarise the GP maps below with detailed comparisons between the three GP maps found in ref. [30].

- *RNA secondary structure*: we use the Vienna package [37] (version 1.8.5) with default parameters to convert RNA sequences to dot-bracket secondary structures. GP maps are represented as RNA*L* with sequences of length *L*.
- *HP lattice model*: we follow refs. [45, 46] and consider energetic interactions between non-adjacent pairs to have values *E*_HH_ = −1, with *E*_HP_ = *E*_PP_ = 0, where *H* are hydrophobic and *P* are polar amino acids. If a sequence has a unique lowest energy structure, its phenotype is that structure, otherwise it is considered degenerate. We consider both the non-compact GP map for all folds of a given length referred to as HP*L* and also only the set of compact structures referred to as HP*l*x*w*x*h*.
- *Polyomino model*: we follow refs. [30, 43] and consider the GP maps 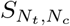 where *N*_*t*_ is the number of assembly kit tiles and *N*_*c*_ with the default self-assembly process used.

The GP maps may be further characterised by their genotype sequence length *L*, base *K*, number of genotypes *N*_*G*_ = *K*^*L*^ and number of phenotypes *N*_*P*_. The redundancy *n*_*p*_ of a given phenotype *p* is the number of genotypes that map to *p* and this is normalised by the size of the genotype space to give the frequency *f*_*p*_ = *n*_*p*_/K^*L*^. The overall redundancy *R* of a GP map is defined as the average number of genotypes per non-deleterious phenotype:

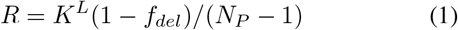

We provide Table III to summarise the characteristic proper-ties used to differentiate the GP maps.

**TABLE III.**
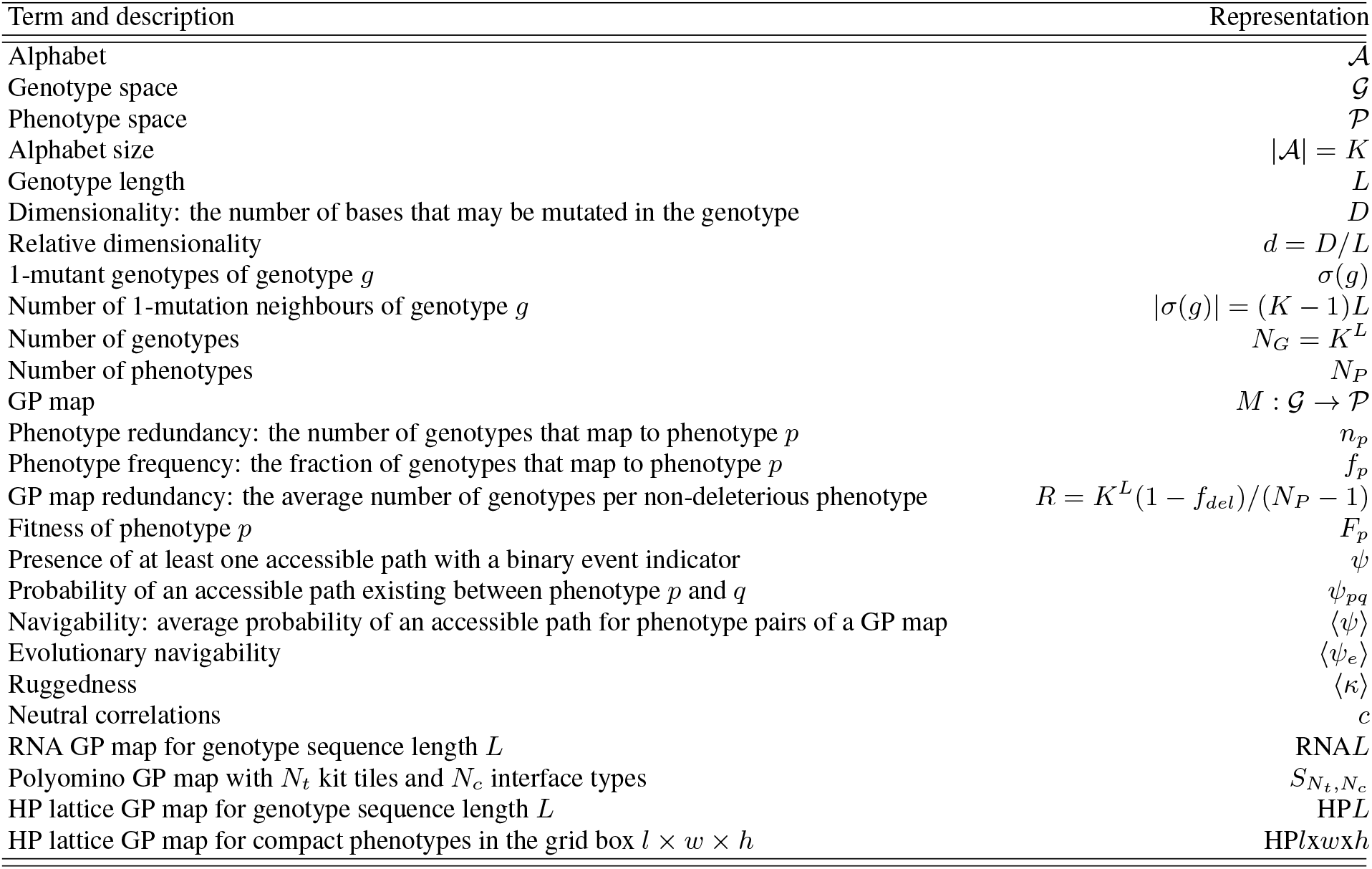
Terminology. A summary of terms and their representations used in the paper.

A particular feature of all three GP maps is a single phenotype that is of a different nature to the others: for RNA secondary structure this is the unfolded ‘trivial’ structure, the HP lattice model it is sequences that have a degenerate ground state and for the Polyomino model it is when there is either unbounded or non-deterministic growth (UND). We refer to this phenotype here as the *deleterious* or *del* phenotype as, in each GP map, we consider it low fitness due to the non-specificity of the structural phenotype. We assign a fitness of zero for *del* throughout this work. While this is a strong assumption, given the large-scale dominance of the *del* phenotype in Polyomino and HP GP maps, we expect this assumption to exacerbate the presence of valleys rather than introducing a bias towards navigability.

### B. Measuring landscape navigability

#### 1. Definitions and formulation

In order to establish the presence of fitness valleys in a fitness landscape, we consider whether it is possible to reach the fittest phenotype from any given point in the genotype space via a path where the fitness increases monotonically defined as an *accessible path* [11, 60]. *Landscape navigability* has previously been defined as the proportion of accessible paths to a given genotype from all other genotypes [17]. To briefly summarise, here we specifically define the navigability as the average probability that a randomly chosen phenotype pair have at least one accessible path between them, given a fitness assignment process to phenotypes. We denote accessibility with *ψ*, where *ψ* = 1 indicates the presence of at least one accessible path between two phenotypes for a specific set of fitness assignments, and *ψ* = 0 indicating no accessible paths. When *ψ* = 0, a fitness valley must be traversed between the phenotypes. With this notation, we use ⟨*ψ*⟩ to represent navigability of fitness landscapes for a given GP map.

#### 2. Fitness landscapes

In conjunction with the GP map *M*, a fitness landscape instance is defined by the set of phenotype fitnesses 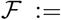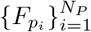, with *i* denoting the *i*^th^ indexed phenotype *p*_*i*_. We refer to the *source* phenotype *p* and *target* phenotype *q* in the search for an accessible path from *p* → *q*. We consider two fitness assignments in this paper:

- *Random fitness*: random samples 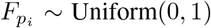 with target phenotype *q* having *F*_*q*_ = 1
- *Hamming distance*: where the similarity of phenotype *p* compared to a phenotype *q* is measured by the number of matching positions in the aligned phenotype string representation given by 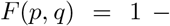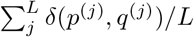, where *p*^(*j*)^ is the string character representing phenotype *p* at the *j*^th^ base position and *F* (*p, q*) is the fitness of phenotype *p* compared to a target phenotype *q*

*F*_*del*_ = 0 for all fitness assignments.

#### 3. Navigability estimation

The probability of an accessible path (*ψ* = 1) between a source phenotype *p* and target phenotype *q*, given a random fitness landscape instance 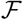, is deterministic with a binary outcome. We can define the probability of *ψ* more explicitly as a function of *p*, *q* and 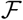 as follows:

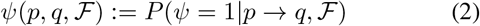

where

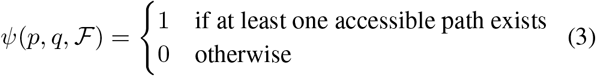

We can take the expectation over 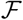 yielding the mean proba-bility of an accessible path from *p* to *q* as:

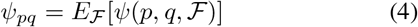

With this notation, we can define t he *navigability* f or the GP map as the expectation over Eq. (4) for phenotypes *p* and *q* sampled uniformly at random:

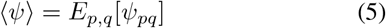

We can estimate this probability of reaching a given target phenotype *q* from a uniform randomly chosen source phenotype *p* by computationally measuring 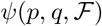 for *N*_*s*_ randomly chosen sources for each of *N*_*t*_ randomly chosen targets, with a new random fitness landscape instance 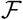 for each pair. We use *I*_*T*_ (*s, t*) to indicate whether the computational estimate for source index *s* with target index *t* was inside the computational threshold *T* and completed the search without aborting. The estimate can be written as:

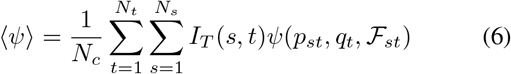

where *p*_*st*_ and *q*_*t*_ are the source and target phenotypes of *s*^th^ source for the *t*^th^ target, the number of completed runs is *N*_*c*_ = Σ_*t,s*_ *I*_*T*_ (*s, t*) and the aborted proportion *α*:

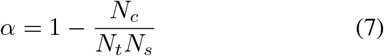

The estimate of the navigability of a fitness landscape with GP map has an associated Bernoulli standard error (derived from an estimate of the corrected sample standard deviation):

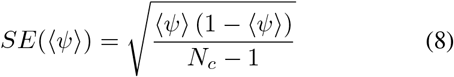

We next describe in more detail the computational algorithm for estimating ⟨*ψ*⟩.

#### 4. Navigability estimation algorithm

For a given source and target phenotype, in each random landscape instance, we perform the following computational algorithm to measure *ψ*. We first provide some definitions:

- GP map *M*: is a function 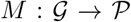 where 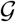 is the space of genotypes and 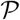 is the space of phenotypes, such that we can write the phenotype *p* of genotype *g* as *p* = *M*(*g*)
- Dimensionality: We define the set of sequence positions that may be mutated as 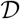, with the size of 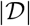 being the dimensionality *D*. When 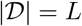 all base positions are mutable. Relative dimensionality is defined as the dimensionality relative to sequence length *d* = *D/L*
- Alphabet: sequences have a set of 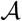 possible letters at a given site. The size of 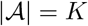 is the base.
- *u*_0_ contains genotypes whose 1-mutant neighbours are yet to be considered in a given search for an accessible path
- *u*_1_ contains genotypes that have already had their 1-mutant neighbours considered in a given search for an accessible path

The algorithm proceeds with a Breadth First Search (BFS):

1. A random genotype *g* that maps to the source phenotype is chosen and added to *u*_0_
2. Set the first element of *u*_0_ as *g*
3. For base 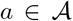 at position *j* and for each position 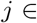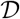, measure genotype neighbour *g*′ and phenotype *p*′ = *M*(*g*′)
4. If *F*_*p*′_ ≥ *F*_*p*_ and *g*′ ∉ *u*_1_, add *g*′ to *u*_0_
5. Move *g* from *u*_0_ to *u*_1_
6. If |*u*_0_| = 0 or |*u*_0_| + |*u*_1_| > *T* (computational thresholdo)r the target phenotype is found, return ‘aborted’ or *ψ* respectively. Otherwise return to step 2

The algorithm finishes with either *u* becoming empty, or the combined size of *u*_0_ and *u*_1_ becoming larger than a predefined threshold *T* (introduced in Section IV B 1), beyond which computational progress may become unfeasible. We discard these aborted runs from the measurement of navigability ⟨*ψ*⟩ using the indicator function *I*_*T*_ of the previous section (Section IV B 3).

As described in Eq. (6) we pick *N*_*s*_ source phenotypes uniformly at random for each of the *N*_*t*_ target phenotypes also chosen at random. We set *N*_*t*_ = 20 and *N*_*s*_ = 50. The uncertainty in the estimate of the navigability ⟨*ψ*⟩ is reported as the standard error *SE*(⟨*ψ*⟩) across the ensemble of measurements.

### C. Removing correlations

In order to measure the effect of positive neutral correlations [30], we perform genotype swaps and then repeat the measurement of ⟨*ψ*⟩. This process involves constructing a new GP map *M*_*s*_ from the original GP map *M*_*s*=0_ ≔ *M* where *s* is the number of pairs of genotypes whose phenotype’s have been swapped. More precisely, a swap involves selecting two genotypes *g*_1_ and *g*_2_ with uniform random probability and setting *M*_*s*_(*g*_1_) = *M*_*s*−1_(*g*_2_) and *M*_*s*_(*g*_2_) = *M*_*s*−1_(*g*_1_). It follows that *M*_*s*→∞_ is the uncorrelated random null model GP map with no positive neutral correlations as used in ref. [30]. As shown in ref. [30], the random null model has *ρ*_*p*_ ≈ *f*_*p*_ when there are no positive neutral correlations. Therefore, we additionally define the correlations *c* present in a given GP map *M*_*s*_ by comparing the logarithm of the average robustness-to-frequency ratio in a given GP map against the original GP map, generating a scale for measuring correlations in *M*_*s*_:

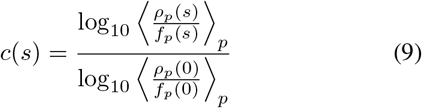

where for *s* = 0 we have *c*(0) = 1, and for lim_*s*→∞_ c(*s*) ≈ 0 the expectation for the random model. Therefore, the scale yields positive values for *c* where there is, on average, greater robustness than frequency. The process of removing correlations gradually from the original GP map (*s* = 0) to the random null model (*s* →∞) provides a range over which the relationship between positive neutral correlations and navigability may be considered in GP maps. We measure the navigability of *S*_2,8_, RNA12, HP3×3×3 and HP5×5 by taking 100 evenly spaced values for *s* on the range *s* = [0, *K*^*L*^] and measuring ⟨*ψ*⟩ and *c*(*s*) for each.

### D. Restricting dimensionality

To measure the role of dimensionality we restrict the dimensionality of a search for an accessible path from source to target by only allowing a set of 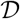 randomly chosen positions along the sequence to be mutated in the 1-mutant neighbour measurement in Step 3 of the navigability algorithm above (Section IV B 4). The dimensionality *D* is the number of positions that may be mutated 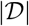, and the relative dimensionality *d* ≔ *D/L*. When *D* = *L* we have the original dimensionality, while for *D* = 1 only a single sequence position may be mutated. The GP map *M* itself is not changed under this dimensional restriction but rather the connectivity of genotypes and therefore the connectivity of the fitness landscape.

We measure the navigability of *S*_2,8_, RNA12, HP3×3×3 and HP5×5 by taking evenly spaced values for *D* on the range *D* ∈ [1, *L*].

### E. Measuring ruggedness

For fitness landscapes, related to navigability is the concept of landscape *ruggedness*. We measure *κ*(*g*), whether a genotype is a local fitness maximum, during the search from source to target. The average proportion of genotypes that are local fitness maxima provides a measure of ruggedness [26]. Whether a genotype *g* is a local fitness peak is determined by the fitness of all accessible 1-mutant neighbours *g*′, such that:

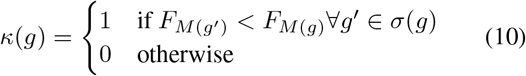

where we have the function *σ*(*g*) which returns the set of 1-mutants of genotype *g*. We calculate the ruggedness for a landscape by taking the average of *κ*(*g*) over all genotypes and all source-target pairs once the search has completed. We denote the ruggedness as ⟨*κ*⟩.

### F. Navigability in the functional RNA database

In Section II C, we examine navigability in a specific subset of RNA phenotypes, namely those that are found in the functional RNA database (fRNAdb) [35]. For a given length we use all phenotypes in proportion to their occurrence in the fRNAdb apart from the trial structure which we exclude as it is assigned zero fitness here. We randomly choose *N*_*t*_ = 50 targets with *N*_*s*_ = 20 randomly chosen sources from this set.

In order to examine navigability between functional RNAs, we must consider sequences longer than *L* = 15. In doing so, we introduce additional computational overhead given the increasing neutral set size resulting in the condition |*u*_0_| + |*u*_1_| > *T* being more likely to be met. Therefore to maximise the number of non-aborted runs, we perform a modified Depth-First Search (DFS) where we attempt to greedily follow paths of increasing gradient until we reach the max fit phenotype. If the path fails, instead of moving back one step as in a standard DFS, we go all the way back to the start of the walk and pick an unexplored neighbour with the lowest fitness to begin a new uphill walk. In this way, we maximise the exploration of new phenotypes by always starting our deep walks from the lowest point while still maintaining the ability to perform long walks during the search.

We write the modified DFS algorithm explicitly as:

1. A random genotype *g* that maps to the source phenotype is chosen and added to *u*_0_.
2. Set the first element of *u*_0_ as *g*, and *p* = *M* (*g*)
3. For each alternative base 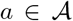 at position *j* and for each position *j* in 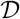, measure genotype neighbour *g*′ and phenotype *p*′ = *M* (*g*′)
4. If any *g*′ has *F*_*p*′_ > *F*_*p*_ and *g*′ ∉ *u*_1_ and *g*′ ∉ *u*_1_, add *g*′ to front of *u*_0_ and return to step 2
5. If any *g*′ have *p* = *p*′ and |*u*_0_| = 1, add one such neutral case to the back of *u*_0_ if *g*′ ∉ *u*_0_ and *g*′ ∉ *u*_1_
6. Move *g* from *u*_0_ to *u*_1_
7. If |*u*_0_| = 0 or |*u*_0_| + |*u*_1_| > *T* (computational thresholdo)r the target phenotype is found, return ‘aborted’ or *ψ* respectively. Otherwise return to step 2.

We note that for searches where neutral mutations are not permitted as part of the search, step 5 of the above is omitted.

### G. Navigability estimation under evolutionary dynamics

We measured fitness landscape navigability as the average probability that a given source-target pair could be connected by way of an accessible path. We extend this definition to the more strict requirement of *evolutionary navigability* where the evolutionary dynamics of a population is considered instead of just the existence of an accessible path in crossing the fitness landscape.

We measure ⟨*ψ*_*e*_⟩ as the proportion of source-target pairs for which the target is reached without the majority population phenotype undergoing a decrease in fitness before finding the target. The majority was taken as being a phenotype that occupied more than 50% of the population’s phenotypes. If no phenotype met this condition in a given generation, then the majority phenotype fitness is not updated.

Evolutionary dynamics were performed using Wright-Fisher dynamics [61, 62], and the additional parameters used for each evolutionary dynamical runs were the following:

- GA parameters: *N*_gen_ = 10, 000, *N*_pop_ = 100, *μ* = 0.05
- fRNA parameters: *L* = 20 and 30

Evolutionary runs that are terminated after *N*_gen_ = 10, 000 generations are treated in the same manner as those that are aborted when estimating ⟨*ψ*⟩. Therefore, evolutionary navigability ⟨*ψ*_*e*_⟩ is the fraction of evolutionary runs that successfully evolved to the target phenotype through the majority population phenotype taking an accessible path across all runs excluding those that were terminated at *N*_gen_ generations.

## V. ACKNOWLEDGEMENTS

The authors would like to thank Marcel Weiß for helpful discussions and insights.

## VI. AUTHOR CONTRIBUTIONS

Conceived and designed the experiments: SFG, AAL, SEA. Performed the experiments: SFG. Analysed the data: SFG, AAL, SEA. Supervised the work: SEA. Wrote the paper: SFG, AAL, SEA.

## Appendix A: Impact of computational thresholds on discovery of estimation of navigability

To allow us to consider the plausibility of navigable landscapes for longer fRNA (*L* > 20), we explore the effect of changing the computational threshold *T* (Section IV B 1) at which the search for an accessible path is aborted. We test four orders of magnitude for the threshold |*u*_0_| + |*u*_1_| < *T* condition: *N*_thresh_ = {2 × 10^3^, 2 × 10^4^, 2 × 10^5^ and 2 × 10^6^}. In each case we attempt *N*_*t*_ = 50 target phenotypes and for each target *N_s_* = 20 source phenotypes and attempt to identify an accessible path, where we record whether a search was successful, unsuccessful or aborted.

In Fig. 5A and Fig. 5B we plot navigability and the proportion of runs that are aborted respectively for the different thresholds against the length of the fRNA sequences. The change in the proportion of aborted runs is pertinent for understanding both how navigability changes when increasing the threshold and also what level of *T* is required to be able to reasonably estimate navigability for a given length *L*. With respect to the first point, in Fig. 5A navigability ⟨*ψ*⟩ ≈ 1 for all lengths *L* and thresholds *T*, showing that almost all non-aborted runs have accessible paths. Extrapolating this observation we should expect high navigability for longer length *L* > 30 if greater computation resource were available. With respect to the required computational thresholds for a given length *L*, we observe, very roughly, that around 50% aborted proportion is reached for *L* = 20 at *T* = 2 × 10^3^, for *L* = 35 at *T* = 2 × 10^5^ and *L* = 40 at *T* = 2 × 10^6^. Extrapolating with quadratic fits we could hypothesise that the aborted threshold could be reduced to 10% for *L* = 40 at between [2 × 10^7^, 2 × 10^8^].

**FIG. 5.**
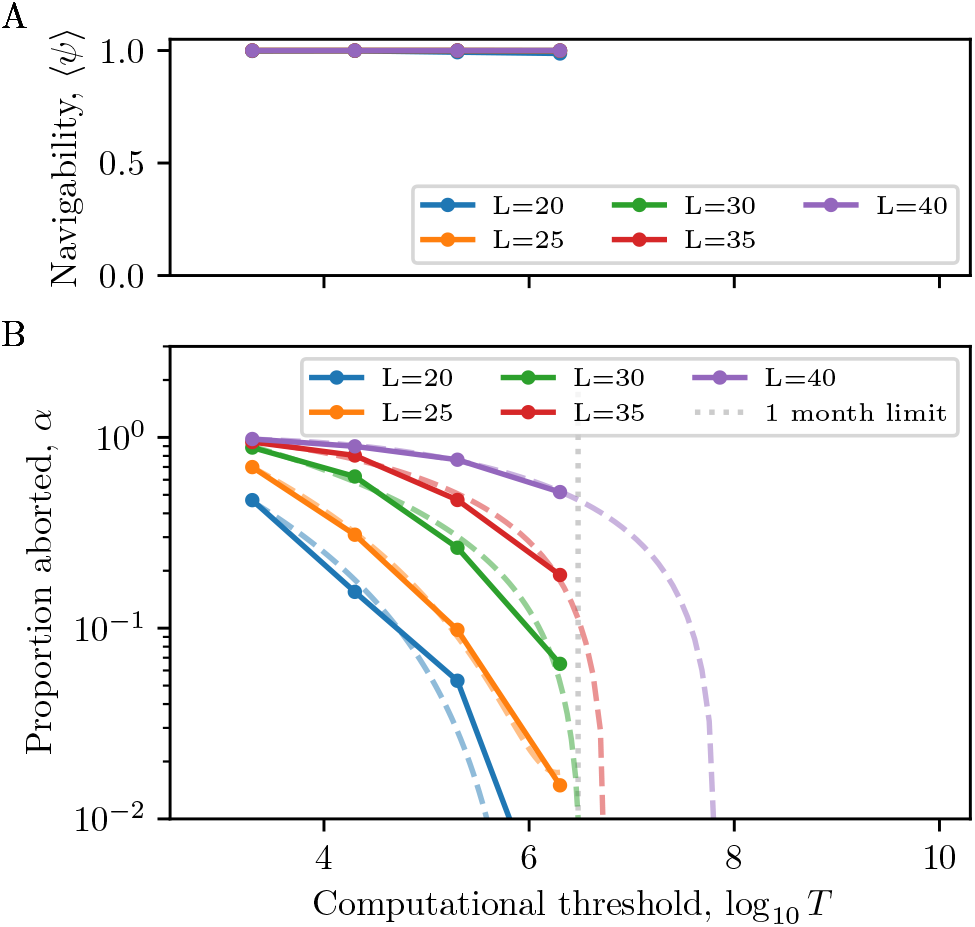
**(A) Navigability ⟨*ψ*⟩ for different length *L* for increasing computational threshold *T*.** Navigability is approximately 1.0 for all computational thresholds suggesting that navigability may be persist for larger computational thresholds. **(B) Proportion of estimations aborted for four different thresholds for different fRNA length *L*.** Dashed lines provide quadratic interpolations to illustrate potential computational thresholds for which a given abortion threshold may be reached if the fit holds for extrapolation. As a guide, we highlight the computational limit corresponding to one month of chronological time given available computational resources.

